# Inflammation-driven cellular selection impacts X-linked mosaicism in the epidermis

**DOI:** 10.64898/2026.10.08.757664

**Authors:** Eleni Angelopoulou, Karan Gandhi, Marjolein van Wolfswinkel, Leonie Schmautz, Inigo Oyarzun, Sarah Nowosadtko, Irmgard Fischer, Viktor Schwartz, Johannes Griss, Stephanie J. Ellis

## Abstract

In eutherian females, the choice of which X chromosome gets embryonically inactivated is thought to be random. Somatic tissues are thus assumed to comprise cells with a 50:50 ratio of maternal:paternal X active. However, a phenomenon known as X-skewing has been observed, wherein tissue composition shifts to be dominated by cells expressing one X or the other. On a gross level, such X-skewing is phenotypically silent except in the case where one X-chromosome carries a disease mutation. Here, by leveraging a mouse model of an X-linked disease called Incontinentia Pigmenti, caused by heterozygous mutations in the NF-kB effector, Nemo, we explore how X-linked cellular dynamics shape progenitor biology in the epidermis. Lineage tracing with X-linked reporters shows that at birth, the epidermis is unexpectedly skewed toward mutant cells. Within a week, as inflammation emerges, these dynamics reverse leaving surviving animals with almost entirely wild-type epithelium. The shift is driven by loss rather than growth since mutant cells adopt an aberrant fate and die at higher rates than their wild-type neighbours; proliferation rises in both populations. Notably, wild-type proliferation scales with local tissue composition, suggesting these cells calibrate their response to the extent of the mutant burden to compensate appropriately for tissue loss. Depleting macrophages, the principal TNF-α source, attenuates inflammation, mutant cell death and ultimately is expected to preserve the mutant population. Altogether, our results identify inflammation as a selective force capable of resolving deleterious mosaicism, not only in the context of an X-linked disease, but within any dynamic, clonally heterogeneous epithelium.

## Results and Discussion

We first sought to verify the previously described phenotype of *Nemo* (gene name: *Ikbkg*) heterozygous mice. Consistent with prior reports^1,2^, we found that Sox2-Cre-driven heterozygous deletion of *Nemo* (hereafter referred to as *Nemo+/-*) resulted in reduced survival compared to wild-type litter mates (**Fig. 1A**), with around 20% of these animals succumbing to the deleterious effects of the genotype. *Nemo+/-* animals exhibited the expected dynamics of epidermal thickening and inflammation starting from around the third postnatal day (P3), peaking at P5, with signs of recovery already appearing at P7 (**Fig. 1B**), and animals at 2 years of age (P609) harbouring epidermal structure indistinguishable from wildtype (**Supplemental Fig. S1A**). Also as previously shown, Sox2-Cre; *Nemo fl/Y* males and *Sox2-Cre;Nemo fl/fl* homozygous females were not viable. These findings already hint at the notion that the presence of some fraction of wild-type cells in the tissue is required to rescue viability. To examine this possibility specifically in the epidermis, we next assessed the post-weaning viability of *Nemo fl/+* versus *Nemo fl/fl* mice harbouring the epidermal specific K14-Cre. Although both genotypes exhibited inflammation emerging early in post-natal life, an overwhelming majority (32/33) of heterozygous *Nemo fl/+* animals fully recovered by weaning at P21 (**Fig. 1C**), also in line with prior reports^3^. In contrast, 100% of *Nemo fl/fl* animals (n = 5) failed to recover. These results are in line with a requirement for epidermal wild-type cells to confer survival.

**Figure 1.**
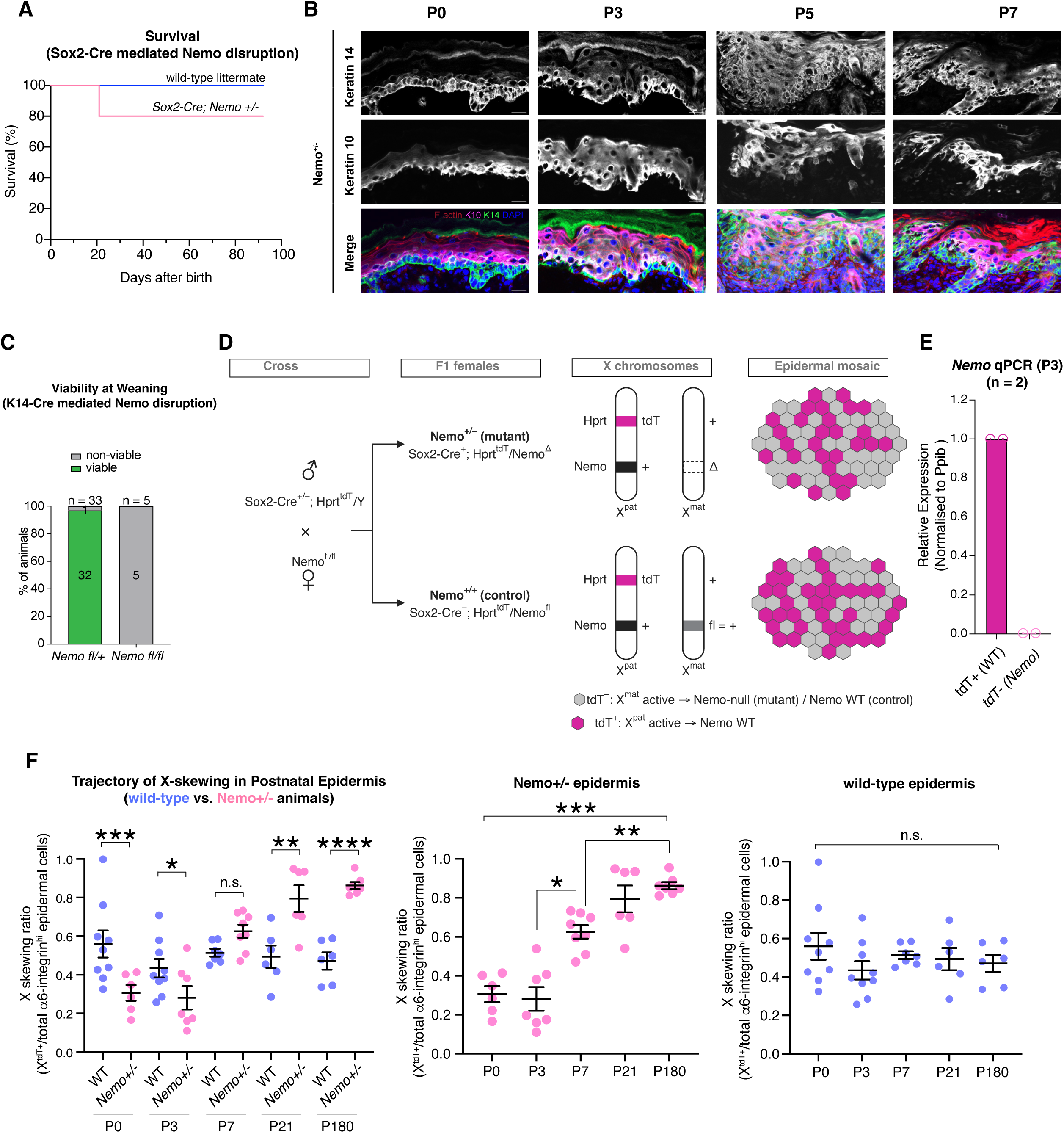
Wild type cells are sufficient to rescue the epidermal phenotype of Nemo +/- females because of X-skewing based cellular dynamics. **(A)** Kaplan–Meier survival of *Sox2-Cre; Nemo* +/– females and wild-type littermates over the first ∼90 days after birth (n = 15 *Nemo +/-*, 15 wild-type). **(B)** Representative immunofluorescence of *Nemo +/-* back skin at P0, P3, P5 and P7 stained for Keratin 14 (K14, basal), Keratin 10 (K10, suprabasal), F-actin and DAPI, showing progressive epidermal thickening and disorganisation between P3 and P5. Scale bars, 20 µm. **(C)** Viability at weaning following epidermis-specific *Nemo* deletion with *K14-Cre*: 32/33 *Nemo fl*/+ and 0/5 *Nemo fl*/fl animals were viable. **(D)** Breeding strategy. *Sox2-Cre*^+^; *Hprt*^tdT^/Y males, which carry a constitutively expressed X-linked tdTomato reporter, were crossed to *Nemo*^fl/fl^ females. All female offspring therefore carry the tdTomato reporter on the paternal X chromosome (X^pat^) and a floxed *Nemo* allele on the maternal X chromosome (X^mat^). In *Sox2-Cre*^+^ offspring, epiblast-wide Cre activity deletes the floxed allele, giving *Nemo*^+/–^ mutant females (*Sox2-Cre*^+^; *Hprt*^tdT/+^/*Nemo^+/^*^Δ^). *Sox2-Cre*^−^ littermates keep the intact floxed allele, which is functionally wild-type, and serve as *Nemo*^+/+^ controls (*Sox2-Cre*^−^; *Hprt*^tdT/+^/*Nemo*^+/fl^). Random X inactivation produces a mosaic epidermis in which tdT+ cells (X^pat^ active) express wild-type *Nemo*, whereas tdT– cells (X^mat^ active) are *Nemo*-null in mutants and *Nemo* wild-type in controls. **(E)** qRT-PCR for *Nemo* on FACS-sorted tdT+ and tdT– epidermal cells from P3 *Nemo +/-* animals, normalised to *Ppib* and expressed relative to tdT+ cells (n = 2 animals). **(F)** X-skewing ratio, measured by flow cytometry and defined as the fraction of tdT+ cells among all alph6-integrin^high^ epidermal cells, in wild-type and *Nemo +/-* epidermis at P0, P3, P7, P21 and P180. Left, comparison between genotypes at each timepoint; middle and right, trajectories within *Nemo +/-* and wild-type epidermis, respectively. In *Nemo +/-* epidermis the wild-type (tdT+) fraction rises from ∼0.3 at birth to ∼0.85 by P180, whereas wild-type controls remain at ∼0.5 throughout. Each dot represents one animal (n = 6–9 animals per group; bars show mean ± SEM). Left: two-way ANOVA (genotype × timepoint) on animal values with Tukey’s multiple-comparisons test. Middle and right: one way ANOVA with Tukey’s post hoc test; selected comparisons shown. Significance: \**P* < 0.05, \*\**P* < 0.01, \*\*\**P* < 0.001, \*\*\*\**P* < 0.0001; n.s., not significant.

In Incontinentia Pigmenti (IP) patients, an absence of a hematological phenotype has been attributed to the observation that blood cells are strongly skewed towards expression of the wild-type allele^4^. To test whether skewing dynamics might also account for survival and phenotype resolution in the epidermis, we took advantage of an X-linked reporter mouse line, expressing the fluorescent protein tdTomato from the *Hprt* promoter^5^. Combining this together with *Nemo +/-* animals and Sox2-Cre (**Fig. 1D**), we first confirmed that tdT+ expression correlated with *Nemo* expression using qPCR; as expected the tdT-population had almost no detectable Nemo signal compared to the tdT+ population in the same animals (**Fig. 1E**). To understand on a population level how skewing patterns tracked with the timeline of the *Nemo +/-* phenotype, we used flow cytometry to follow the expression of the tdT+ wild-type allele versus the expression of the unmarked tdT-*Nemo* mutant allele in single cell suspensions from the epidermal fraction of early postnatal backskin. We then carried out a time course analysis to interrogate the proportion of tdT+ within the epidermis from P0, through to P180 (**Fig. 1F**). This analysis revealed that at P0, there was greater representation of tdT+ cells in wildtype epidermis than in *Nemo+/-* epidermis where more of the suspension was tdT-. This suggested that already from birth and persisting at P3, *Nemo* mutant-expressing cells have an advantage compared to their wild-type counterparts. However, as the inflammatory phenotype progressed, we observed a gradual swing in the population dynamics such that by P7 more than half of the epidermal cells were tdT+. By P180, nearly all of the epidermis was comprised by tdT+ wild-type cells, indicating that the tdT-*Nemo*-mutant expressing allele population was largely eliminated.

Our observations raise the question of what cellular dynamics underlie this dramatic, population level change in tissue composition. First, we considered apoptosis using cleaved caspase-3 (CC3) staining. This revealed an increase in signal at the tissue level that peaked at P5 (**Fig. 2A-B**), before already beginning to resolve at P7, although this decline did not reach statistical significance (**Fig. 2B’**). By leveraging the power of our X-linked reporter system, we could specifically attribute the increase in apoptotic cell death to the tdT-Nemo mutant cell population starting from P3 (**Fig. 2D**). In contrast, the wild-type tdT+ population exhibited no increase in caspase-3 staining compared to wild-type controls (**Fig. 2C**). Interestingly, although we did not see a statistically significant increase in caspase-3 activity at the tissue level at P0, by performing paired comparisons of tdT+ versus tdT-cells within the same fields of view, we discovered a small but significant increase in caspase-3 signal at P0, and between the two populations at all subsequent time points (**Fig. 2E**). Taken together, we conclude that apoptotic cell loss is a major driver of Nemo mutant cell loss.

**Figure 2.**
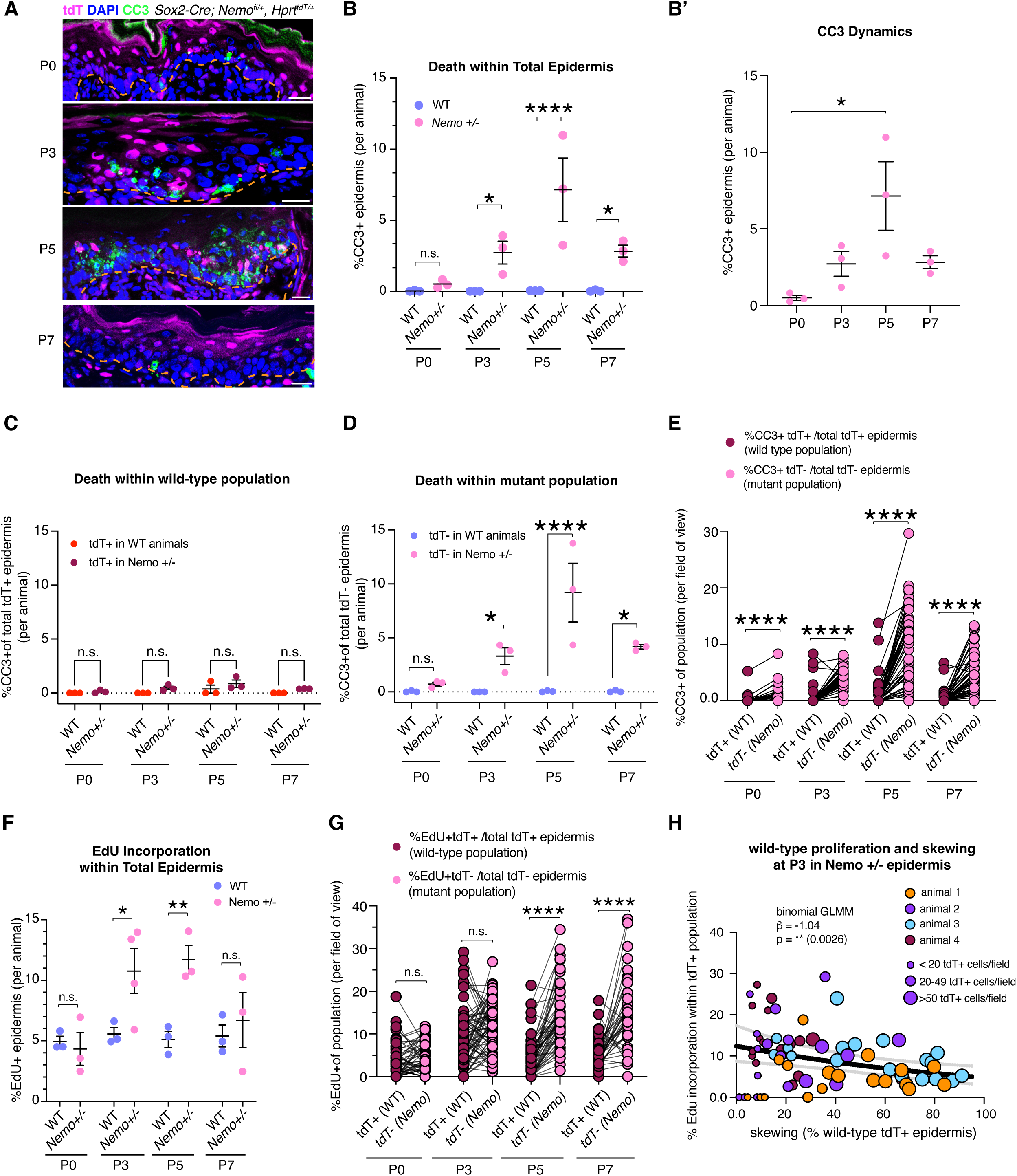
Mutant cell death is the primary driver of population-level skewing. **(A)** Representative images of *Sox2-Cre; Nemo fl*/+; *Hprt*tdT/+ epidermis at P0, P3, P5 and P7 stained for cleaved caspase-3 (CC3), with tdTomato and DAPI. Dashed line marks the basement membrane. Scale bars, 20 µm. **(B)** Percentage of CC3+ cells in the total epidermis of wild-type (*Nemo*+/+) and *Nemo +/-* animals. **(B′)** Data for *Nemo +/-* animals from (B) replotted over time; apoptosis peaks at P5. **(C, D)** Percentage of CC3+ cells within the tdT+ (C) and tdT– (D) epidermal populations of wild-type and *Nemo +/-* animals. Apoptosis is selectively increased in the tdT– (*Nemo*-null) population of *Nemo +/-* animals from P3 onward. **(E)** Within-field comparison of the percentage of CC3+ cells in the tdT+ (wild-type) and tdT– (mutant) populations of *Nemo +/-* epidermis; lines connect populations from the same field of view. **(F)** Percentage of EdU+ cells in the total epidermis of wild-type and *Nemo +/-* animals (1h EdU pulse). **(G)** Within-field comparison of EdU incorporation in tdT+ and tdT– populations of *Nemo +/-* epidermis, as in (E). **(H)** EdU incorporation in the tdT+ population as a function of local skewing (% tdT+ basal cells) per field of view in P3 *Nemo +/-* epidermis. Each point is one field; colour indicates animal (n = 4 animals) and point size the number of tdT+ cells per field. Line shows a binomial generalised linear mixed model fit (response, EdU⁺ of tdT⁺ cells per field; fixed effect of local tdT⁺ fraction; random intercepts for animal and for field, the latter accommodating overdispersion; n = 71 fields from 4 animals), with 95% confidence band. Odds of EdU incorporation decrease by ∼10% per 10-percentage-point increase in local tdT⁺ fraction (odds ratio 0.90, 95% CI 0.84–0.96; β = −1.04 log-odds per unit fraction, P = 0.0026). (B–D, F) Each dot represents one animal (n at least 3 per genotype and timepoint); bars show mean ± SEM; two-way ANOVA (genotype x timepoint) on animal means followed by uncorrected Fisher’s LSD tests comparing genotypes at each timepoint. (B′) one-way ANOVA with Tukey’s test for multiple comparisons. (E, G) Each dot represents one field of view from n = 3 animals per timepoint; paired Wilcoxon signed-rank test at the field level. Significance: \**P* < 0.05, \*\**P* < 0.01, \*\*\**P* < 0.001, \*\*\*\**P* < 0.0001; n.s., not significant.

We next explored whether proliferative dynamics also tracked with overall skewing dynamics by measuring EdU incorporation as a readout for the percentage of cells in S-phase. Consistent with an inflammatory response, we observed an overall increase in the proportion of EdU+ epidermal cells compared to wild-type controls at P3 across the entire epidermis. This difference persisted at P5, but was no longer apparent at P7, consistent with the resolution of the inflammatory phenotype that starts to occur around this time point (**Fig. 2F**). Further stratifying these dynamics by mutant (tdT-) versus wildtype (tdT+) population, we asked whether mutant cells might have a relative deficit in proliferative capacity compared to wild-type neighbours. However, we found that both populations contributed to the overall increase in EdU+ epidermis in a timepoint-dependent manner, with the wild-type having a significant increase at P3 and the mutant population significant later at P5 (**Supplemental Fig. S1B, C**). Interestingly, upon performing paired comparisons, we found there was no difference in EdU incorporation between mutant and wild-type cells at P3, but that at P5 and P7, the mutant cells were incorporating more EdU than wild-type cells, an observation standing somewhat in contrast to the population-level changes in representation we observed (**Fig. 2G**). Although it is challenging to infer quantitative death and proliferation dynamics using the tools at our disposal, we conclude that the increase in mutant cell proliferation is not sufficient to compensate for the massive increase in death observed. On the other hand, the wild-type cells, despite displaying elevated proliferation for a shorter time window, experience far less loss and thus are able to compensate and gradually replace the dying Nemo mutant population. To more quantitatively explore the possibility that wild-type cells tune their response according to tissue composition, we performed a correlation analysis on imaging data from single fields of view examining the relationship between local skewing in terms of the percentage of wild-type epidermal cells, and the percentage of tdT+ cells that were EdU+ within that same region. This approach revealed a negative correlation between the skewing towards wild-type cells and wild-type EdU incorporation suggesting that within local regions of the epidermis, wild-type proliferation scales inversely with the local proportion of wild-type cells, consistent with a compensatory response to mutant cell burden (**Fig. 2H**). Critically, performing the same analysis on the tdT-mutant cell population, we did not observe a statistically significant relationship (**Supplemental Fig. S1D**).

We next sought to dissect the molecular underpinnings of the cellular behaviors we observed across the timeline of the change in skewing dynamics. We thus performed single-cell RNA-sequencing (scRNA-seq) of single-cell suspensions on both *Nemo +/-* and wild-type epidermis. This approach uncovered 12 distinct clusters across three time-points that largely corresponded to the expected cell populations in the epidermal fraction of the skin, including stromal cell types (**Fig. 3A-B, Supplemental Fig. S2A**). However, two of these clusters were present exclusively in the *Nemo +/-* epidermis: a “wound” cluster and a “stressed keratinocyte” cluster that were named so because of their highly abundant expression of specific keratins associated with repair and/or general epidermal pathology (**Supplemental Fig. S2B-D**). These include *Krt16*, *Krt6a/b*, and *Krt17*. These clusters tracked the inflammatory phenotype: beginning to emerge at P0, becoming significantly pronounced at P3, and regressing by P7. An important confound to this observation, however, is that the P7 data in both Nemo +/- and wild-type animals was dominated by hair follicle lineages that are in full anagen, and thus the overall representation of epidermal cell types in those samples is comparatively less than the other two time points. Pseudotime analysis suggested the wound and stress clusters represented an aberrant population, diverted from the normal path of epidermal maturation (**Supplemental Fig. S2E**).

**Figure 3.**
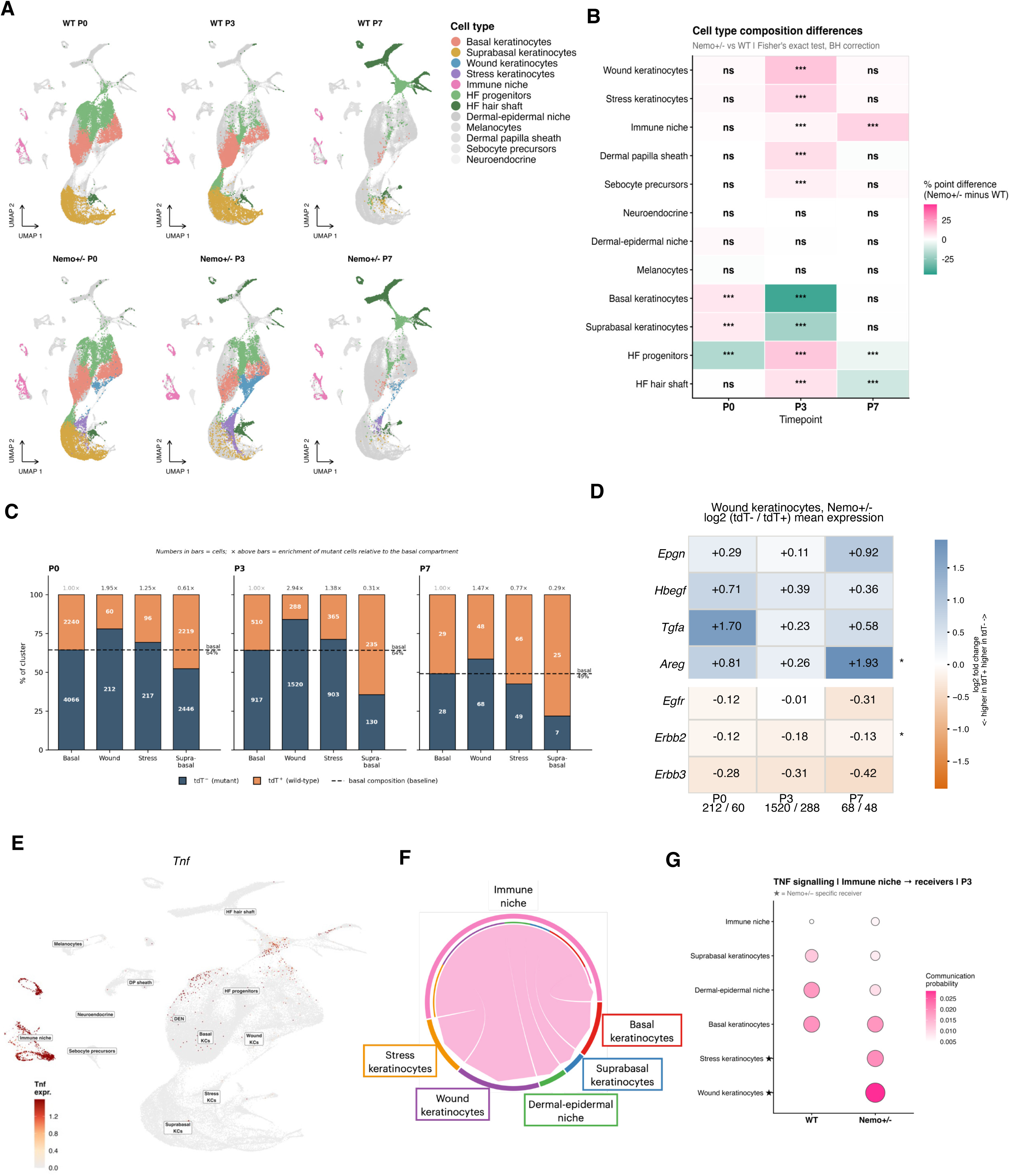
scRNA-seq reveals a mutant-enriched, wound-like keratinocyte population that is hyper-responsive to immune-derived TNF-α. **(A)** UMAPs of the MNN-integrated single-cell RNA-seq dataset from wild-type and *Nemo +/-* back skin at P0, P3 and P7 (n = 2 animals per genotype and timepoint), coloured by annotated cell type; cells from the indicated sample are coloured and all other cells are shown in grey. Cluster markers are shown in Supplementary Figure 3. **(B)** Differences in cell-type composition between *Nemo +/-* and wild-type samples (percentage-point difference, *Nemo +/-* minus wild-type) at each timepoint. Fisher’s exact test with Benjamini– Hochberg correction. ***adjusted *P* < 0.001; ns, not significant. **(C)** Proportion of tdT– (mutant) and tdT+ (wild-type) cells in basal, wound, stress and suprabasal keratinocyte clusters of *Nemo +/-* epidermis at P0, P3 and P7. Numbers within bars indicate cell counts; values above bars give the odds of a cell being tdT– relative to the basal keratinocyte compartment (dashed line, basal composition). Wound keratinocytes are enriched for mutant cells at P0 (1.95×) and P3 (2.94×). **(D)** Heatmap of log2 fold change (tdT– / tdT+) in mean expression of EGF-family ligands (*Epgn*, *Hbegf*, *Tgfa*, *Areg*) and receptors (*Egfr*, *Erbb2*, *Erbb3*) in wound keratinocytes of *Nemo +/-* epidermis. Numbers below columns give tdT– / tdT+ cell numbers per timepoint. *adjusted *P* < 0.05, pseudobulk DESeq2, P0–P7 pooled (see Supplementary Fig. 4C). **(E)** Feature plot of *Tnf* expression on the integrated UMAP. **(F)** CellChat chord diagram of inferred TNF signalling from the immune niche to epidermal populations in P3 *Nemo +/-* skin. **(G)** CellChat communication probability for TNF signalling from the immune niche to receiver populations at P3 in wild-type and *Nemo +/-* skin. Stars mark receivers detected only in *Nemo +/-* samples (stress and wound keratinocytes).

We next asked, using our tdT reporter system, where the mutant versus wild-type cells fall within the clusters we uncovered. At P0 and P3, consistent with the overall pattern of skewing in the tissue, the basal cell cluster was dominated by mutant cells (**Fig. 3C**). Moreover, the wound cluster, and to a lesser degree the stressed keratinocyte cluster was also majority mutant cells. The fact that we found some wild-type cells also resided in these populations suggested at least in part that these aberrant states are not genotype specific but correspond to the tissue-level disease program. On the other hand, the differentiated compartment of the skin was largely dominated by wild-type cells, consistent with the impression from our other analyses that mutant keratinocytes more often entered aberrant wound-like cell states. Differential expression analysis of mutant versus wild-type cells in the wound cluster revealed few differences in gene expression between genotypes, although consistent with their respective position in the tissue, mutant cells tended to have higher expression of basement membrane and integrin signaling genes whereas wild-type cells were enriched in epidermal differentiation pathways (**Supplemental Fig. S2F-G**). Sample preparation for RNA-sequencing largely excludes dying and dead cells, but the mutant-biased composition and timing of the emergence of the wound cluster suggest it may represent the population that subsequently undergoes apoptosis.

Our analysis of cellular behavior raised the question of whether there was crosstalk between mutant (tdT-) and wild-type (tdT+) epidermal populations that enables the wild-type cells to compensate for lost mutant cells. To address this prospective mechanism, we turned to CellChat ^6^, an algorithm that infers cell-cell interactions from expression of ligand-receptor pairs between cell populations. Among the interactions it identified, EGF-family signaling offered a direct and testable explanation for the potential non-autonomous proliferative response we observed in wild-type cells. We therefore examined EGFR specifically.

Focusing on the wound cluster, pseudobulk differential expression analysis showed that mutant cells express higher levels of the EGFR ligand *Areg* (padj = 0.019), with the remaining EGF ligands showing the same direction without reaching significance (**Fig. 3D, Supplemental Fig. 4A,C**). Commensurate with this, wild-type cells within the same cluster expressed higher levels of the EGFR co-receptor *Erbb2* (padj = 0.019), with *Egfr* itself as well as the other co-receptor *Erbb3* trending in the same direction (**Fig. 3D, Supplemental Fig. 4B,C**). Importantly neither the basal nor the stress keratinocyte clusters showed any such differences (all padj>0.9), suggesting that this signaling asymmetry is specific to cells that have entered the aberrant wound state rather than a general feature of the mosaic epidermis. These observations raise the possibility, as observed in injured homeostatic mouse skin^7^ and the Drosophila gut^8^, that mutant cells within the wound state, which we suspect represent the population destined to die, secrete mitogens to promote a compensatory proliferative response from non-dying neighbours. This yet untested mechanism would facilitate both expansion of the wild-type cell population and eventual restoration of homeostasis.

A second implication of our cellular analyses is that mutant cell apoptosis is the critical cue that triggers that directional shift in X-skewing dynamics. But what is the molecular trigger? Prior literature predicts that macrophage TNF-α is the stimulus that leads to *Nemo* mutant cell apoptosis since *Nemo* keratinocytes in culture are sensitive and die upon TNF-α treatment and TNF-α is only present in the tissue from around post-natal day 3; the inflammatory phenotype of *Nemo* null mice has also been shown to be TNFR1 dependent^3^. More recent work, as well as our own data (**Fig. 3E**), suggested that macrophages are a major source of TNF-α in postnatal skin, signaling via TNFR1 on epidermal stem cells to promote clonal expansion^9^. To probe this in our dataset we again used CellChat to ask which cell populations were the predicted source and targets of TNF signalling. As expected, this identified the immune cell cluster as the dominant sender in both wild-type and Nemo +/- animals (**Fig. 3F**). Although all epidermal clusters in both genotypes were predicted receivers, within the Nemo +/- animals, the wound and stress clusters had the greatest receiver strength (**Fig. 3G**). We also observed a small but statistically significant increase in the immune cell cluster at P3, a finding that correlated well with dermal infiltration of macrophages also observed around this time point (**Supplemental Fig. S4D**).

These analyses together predicted that myeloid immune cells (likely macrophages) are the dominant source of TNF-α in *Nemo +/-* skin, and that the aberrant keratinocyte populations are among its principal targets. Together with the established TNFR1-dependence of the epidermal phenotype, this suggested that immune cell-derived TNF-α drives the elimination of mutant cells and, consequently, the shift in skewing dynamics. To test this directly, we depleted Csf1r-dependent myeloid cells using custom chow formulated with the Csf1r inhibitor PLX5622; this approach has been used previously to deplete myeloid cells from the periphery including the developing skin^10–12^. *Nemo +/-* pups born to mothers who had been on the chow since E10.5 showed significant reduction of F4/80+ macrophages infiltrating the dermis at P5 compared to *Nemo+/-* pups on the control diet (**Fig. 4A-B**). Moreover, *Nemo+/-* pups displayed a striking reduction in epidermal thickness compared to pups on the control diet (**Fig. 4C**), and apoptosis was markedly reduced (**Fig. 4D**). Critically, few tdT+ cells were visible within the basal layer at P5, suggesting that skewing towards wild-type cells was not underway (**Fig. 4E**). Taken together, these data place macrophage infiltration and subsequent apoptosis as the initiating force for mutant cell elimination. We speculate they are the master orchestrator of the dramatic X-skewing dynamics we observe.

**Figure 4.**
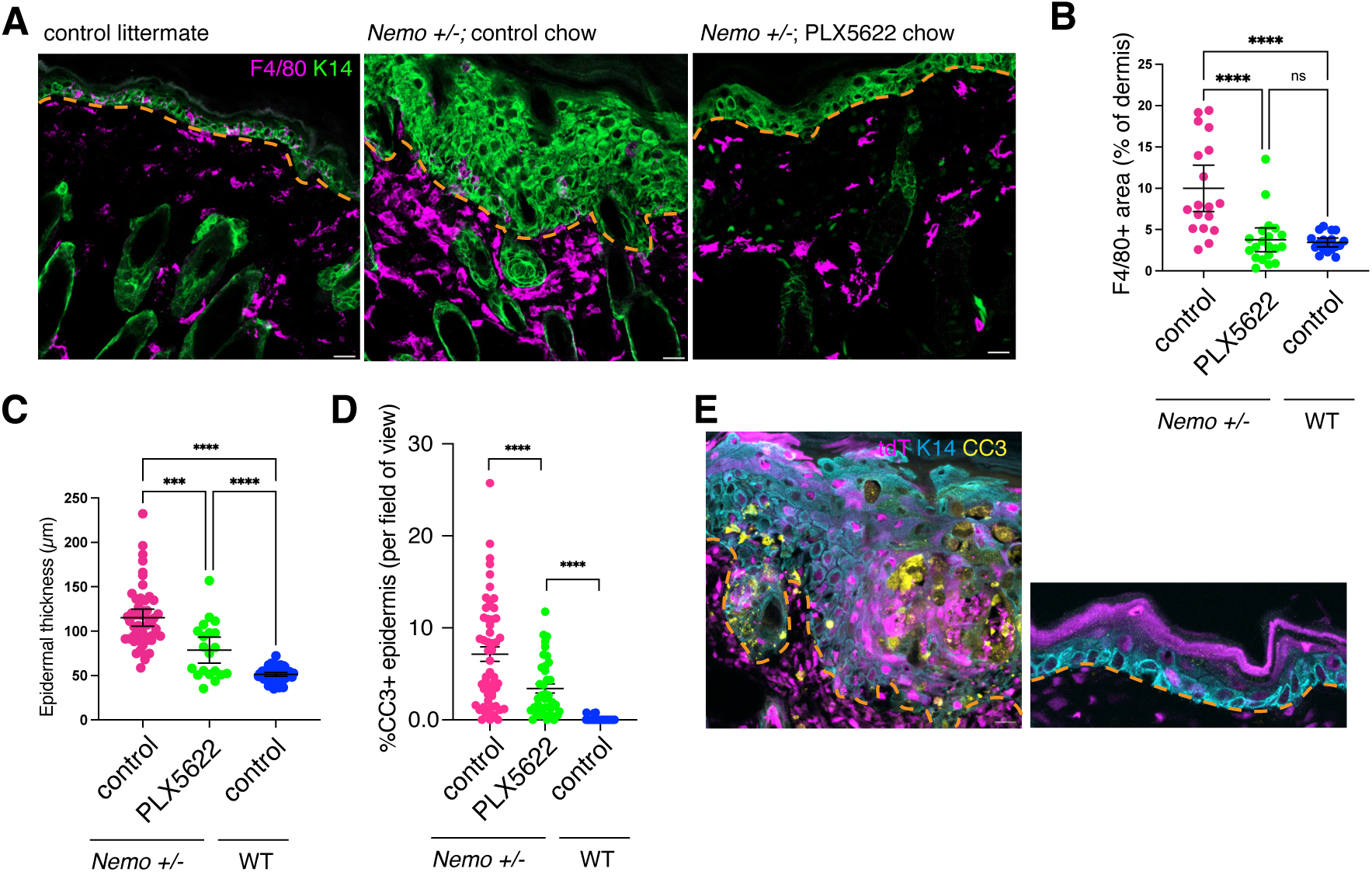
Macrophage depletion attenuates the inflammatory phenotype and mutant cell apoptosis in *Nemo +/-* epidermis. **(A)** Representative images of back skin from a wild-type littermate and *Nemo +/-* animals fed control chow or the CSF1R inhibitor PLX5622 from E10.5 (via the dam), stained for F4/80 and K14 at P5. Dashed line marks the basement membrane. Scale bars, 20 µm. **(B)** Dermal F4/80+ area (% of dermis). **(C)** Epidermal thickness. **(D)** Percentage of CC3+ cells in the epidermis. **(E)** Representative images of tdTomato, K14 and CC3 in *Nemo +/-* control chow (left) and PLX5622 (right) skin. Scale bars, 20 µm. (B–D) Each dot represents one field of view from n = at least 2 animals per group; bars show mean ± SEM; one-way ANOVA. Significance: \**P* < 0.05, \*\**P* < 0.01, \*\*\**P* < 0.001, \*\*\*\**P* < 0.0001; n.s., not significant.

Disease phenotypes in heterozygous female patients of X-linked diseases are known to be a product of the balance of wild-type versus disease allele expressing cells. However, the tools and technology to understand what sets this ratio, and to map how wild-type and mutant cells interact with one another have only become available recently. In the absence of these tools over twenty years ago, it was posited that mutant cells within the IP mouse model were subject to cell competition-like selection during disease resolution^3^. Here we show that wild-type cells do not outcompete mutant cells, but rather compensate for the tissue lost to cell-autonomous mutant cell apoptosis, driving disease resolution. Mutant keratinocytes are intrinsically sensitised to TNF-induced death and die even in a fully mutant epidermis^3^, and we find no evidence that their loss depends on the local composition of the surrounding tissue — the fitness sensing between neighbours that defines competition^13^. This stands in contrast to recent work in the hematopoietic system, where analysis of heterozygous *Stag2* mutations revealed that the continued presence of wild-type cells is required to exclude mutant clones from the lymphoid lineage^14^. In our case wild-type cells respond to mutant cell loss, whereas in the *Stag2* example they provide the initiating force for population removal.

More broadly, our findings position inflammation as a selective filter that shapes clonal dynamics and disease outcomes depending on how a given mosaic mutation alters tolerance to inflammatory signals. For example, in the haematopoietic system, inflammation favours the expansion of tolerant mutant clones that drive clonal hematopoeisis^15–17^. Similarly, in the epidermis, immune-cell derived TNF-α promotes clonal expansion of pre-malignant mutations ^9^. Intriguingly, in our case, we find the opposite: that an overly-sensitized mutant population is eliminated by inflammatory signals. For IP, this suggests that the inflammatory phase, though pathological, may be what ultimately resolves the disease state. Therefore, although an otherwise attractive therapeutic avenue, dampening this response could carry the cost of preserving mutant cells, which would leave the tissue highly vulnerable to later-in-life insults that trigger inflammation, and perhaps even tumorigenesis.

## Acknowledgements

We thank M. Pasparakis for the floxed Nemo allele. The scRNA-sequencing (including RNA isolation and library preparation) was performed by the Next Generation Sequencing Facility at Vienna BioCenter Core Facilities (VBCF), member of the Vienna BioCenter (VBC), Austria. We thank Max Perutz Labs BioOptics FACS Facility for flow cytometry and sorting services and the Max Perutz Labs Light Microscopy Facility microscopy support. We thank A. Khandekar for designing the probes for the RNA-sequencing, and S. Sharma for technical assistance. We thank all members of the Ellis lab for valuable discussions. This work was supported by an FWF-START award (to S.J.E.) and a Vallee Foundation Scholar Award (to S.J.E.).

## Experimental Procedures

### Mouse lines and animal survival

#### Mouse lines and genotyping

Mouse breeding and experimental protocols performed during this project were authorised by the Austrian Ministry (BMFWF). All mice were housed under specific pathogen-free conditions in the MPL animal facility. No statistical methods were used to predetermine sample size.

Most experiments were performed using *Sox2-Cre^tg/wt^;Hprt^tdTomato/wt^,Nemo^Δ/+^* females and wild-type (*Sox2-Cre^wt/wt^;Hprt^tdTomato/wt^;Nem^/fl/+^*) littermates in a CD1 mixed background. This line was generated by first crossing *Sox2-Cre^tg/wt^* (JAX stock #008454) males^18^ with *Hprt^lox-stop-lox-tdTomato/lox-stop-lox-tdTomato^*(JAX stock #021428)^5^ females. This generated males that constitutively express the *Hprt*-linked tdT reporter, including in the germline (*Hprt^tdTomato/Y^; Sox2-Cre^tg/wt^*). These males were then crossed to *Nemo^fl/fl^* females^2^ to generate the required genotypes for experiments.

Additionally, *K14-Cre^tg^;Nemo fl* females and males were generated by first crossing *K14-Cre^tg^* (JAX #018964)^19^ females with *Nemo fl^/Y^* males. Then, *K14-Cre;Nemo fl^/+^* females were crossed with *Nemo fl^/Y^* males. *K14-Cre;Nemo fl^/fl^* females, *K14-Cre;Nemo fl^/+^* females, and *K14-Cre;Nemo fl^/Y^* males were used for experiments. Mice were genotyped using the following primers:

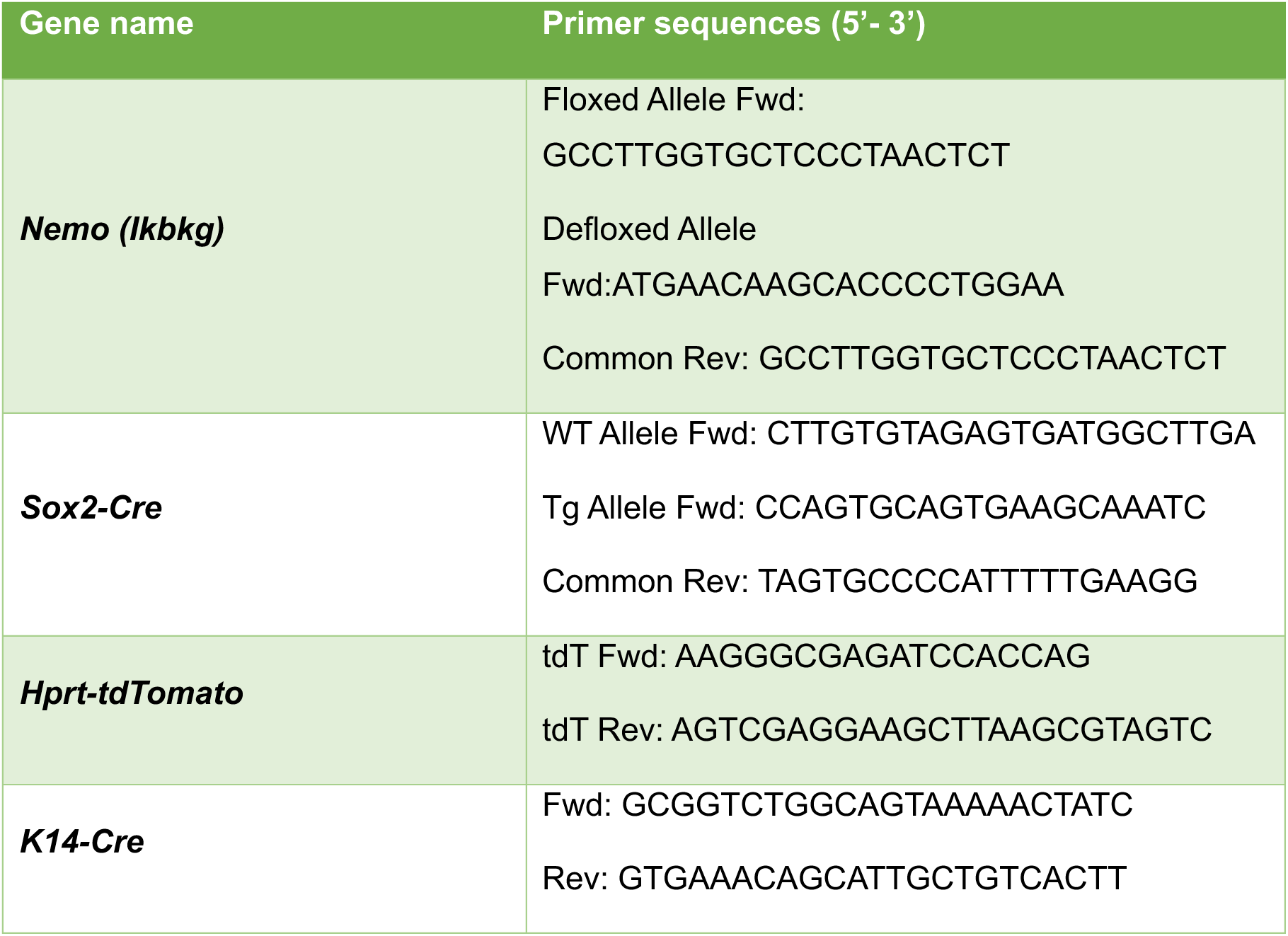

#### X-reporter system

The Hprt-tdTomato (Hprt-tdT) reporter is a transgenic allele at the X-linked Hprt locus, such that cells in which the Hprt-tdT X chromosome is active express tdTomato fluorescence. In *Nemo +/−* animals, the *Hprt-tdT* allele was carried on the same X chromosome as the wildtype *Nemo* allele, such that tdT-positive cells express and tdT-negative do not. In wild-type animals, tdT-positive cells represent cells where the paternal X is active, with no selective pressure between the two populations expected as previously reported in Wu *et al* ^5^.

#### Animal survival analysis

Survival curves were plotted using data from *Sox2-Cre^tg/wt^;Hprt^tdTomato/wt^,Nemo^Δ/+^* female pups (and corresponding wild-type controls) or K14-Cre; *Nemo* pups. In the first case, survival was recorded until postnatal day 92. Animals found unalive were recorded as death events. Monitoring the survival of *K14-Cre;Nemo^fl/fl^* females, *K14-Cre;Nemo^fl/+^*females, and *K14-Cre;Nemo^fl/Y^* males lasted from birth until weaning (day 21), since severely affected animals were not expected to survive beyond weaning.

Survival curves were plotted using GraphPad Prism (version 11.0.0) and the Kaplan-Meier method, comparing between genotypes using the log-rank (Mantel-cox) test. Hazard ratios and 95% confidence intervals were calculated using the Mantel-Haenszel method, with p-values below 0.05 were considered statistically significant.

### Macrophage depletion

To deplete macrophages from *Sox2-Cre^tg/wt^;Hprt^tdTomato/wt^,Nemo^Δ/+^* pups, *Nemo^fl/fl^* females were crossed to *Hprt-tdT^tdTomato/Y^; Sox2-Cre^tg/wt^* males, and were separated after a positive plug (day E0.5). On day E10.5, standard chow was replaced by custom chow formulated with the selective Csf1r-inhibitor PLX5622 at a dose of 1200ppm (drug purchased from TargetMol and formulation done by Research Diets, Inc.), and was available ad libitum until pups were sacrificed. Back skin samples were obtained from pups at P5.

### qPCR

Back skin was collected from experimental mice, and single cell suspensions were made. Single cells were sorted using the Aurora CS Cell Sorter into tdT-positive and tdT-negative. The sorted cells were transferred to TRIzol and RNA was isolated using the Monarch Total RNA Miniprep Kit as per manufacturer protocol. cDNA was synthesized using the LunaScript RT SuperMix Kit. qPCR was carried out using the Luna Universal qPCR Mastermix. Cq values for the target genes were normalized to Ppib, which served as a control. The data was analysed using the Delta Delta Ct Method. The primers are sequences (5’ to 3’) are as follows: *Nemo (Ikbkg):* Fwd: TGTGCAAATTCCAGGAAGCC, Rev: GAGTTCTCCGAGCAATGATGTC, with a product size of 179 bp, and *Ppib:* Fwd: GTGAGCGCTTCCCAGATGAGA, Rev: TGCCGGAGTCGACAATGATG, with a product size of 252 bp.

### Histology

Back skin was dissected from mouse pups at P0, P3, P5, and P7. The skin was fixed in 4% (w/v) Paraformaldehyde (Fisher Scientific, 11400580) for 30 min at 4 °C and washed in 1X PBS three times on an orbital shaker. The fixed tissue was incubated in 30% Sucrose overnight at 4 °C, after which it was cut in strips and embedded in O.C.T. Compound (Tissue-Tek, 4583) and placed at -80 °C overnight. Frozen tissue was cut in 12-14 μm thick sections using a Cryostat (Epredia, NX70) and kept at -20 °C.

### Immunofluorescence and microscopy

Frozen sections were prepared for staining by washing in 1X PBS, permeabilising with 0.3% Triton X-100 PBS (PBS-T), and blocking in 0.3% blocking buffer (2.5% fish gelatin, 5% donkey serum, 1% bovine serum albumin, in 0.3% PBS-T). Slides were incubated overnight at 4 °C with primary antibody mix in 0.3% blocking buffer. Primary antibodies used were as follows: Guinea pig anti-cytokeratin 14 (Antibodies Online, ABIN113455, 1:500 dilution), Rat anti-RFP (Chromotek, 5F8, 1:200 dilution), Rabbit anti-RFP (MBL, PM005, 1:200 dilution), Rabbit anti-Cleaved-Caspase3 (Cell Signalling, 9661, 1:200 dilution), Rat biotinylated anti-CD45 (Biolegend, 5530, 1:200 dilution), Rat anti-F4/80 (Invitrogen, MF48000, 1:200 dilution). After three 0.3% PBS-T washes, slides were incubated with secondary antibody mix in 0.3% blocking buffer at RT for 2.5 h. Secondary antibodies were raised in donkey host and conjugated in AlexaFluor488, Cy3, or AlexaFluor647 (Jackson ImmunoResearch Lab, 1:500 dilution). After incubation, slides were washed in 0.3% PBS-T and 4’,6-diamidino-2-phenylindole (DAPI, 1:5000 dilution) was used at RT for 15 min to label nuclei. After staining, slides were dried and mounted using ProLongTM Diamond Antifade Mountant (Molecular Probes, P36961) and stored at 4 °C until imaging.

Sections were imaged using the 40x oil objective on a ZEISS LSM980 confocal microscope, operated by ZEN microscopy software. Six fields of view from three skin sections each, for three animals per genotype were imaged and analysed.

### EdU labelling

Pups were weighed and subcutaneously injected (25 μg/g) with 5-ethynyl-2′-deoxyuridine (EdU, Invitrogen, A10044). They were sacrificed 1 h post injection and the back skin was collected and prepared for staining as described earlier. EdU labelling was performed using the Click-iT Alexa Fluor 647 Imaging kit (Invitrogen, C10340), following the manufacturer’s instructions. Briefly, after incubation with primary antibody mix and subsequent washes, 130 μl of the EdU cocktail mix were added per slide and slides were incubated at RT in the dark for 45 min. Then they were washed in 0.3% PBS-T and antibody staining was performed as previously described.

### Image analysis

Image analysis was performed using Fiji (ImageJ2, version 2.16.0/1.54p^20–22^). For every image, the epidermal region of interest was defined manually using cytokeratin 14 (K14) immunofluorescence as a basal layer marker, and the epidermis was defined as the tissue above the K14-positive layer. Cell counting was performed manually using the Fiji Cell Counter plugin. For each analysis, a combination of the following populations was counted: total tdT-positive and tdT-negative cells, EdU-positive cells, CC3-positive cells, and CD45+ (or F4/80+) cells, with each marker assessed within the defined epidermal region.

Quantification of the dermal CD45+ region (or F4/80+ in the case of Fig. 4), the CD45-positive area fraction within the dermis (below the K14 boundary) was measured using thresholding in Fiji. The X-skewing ratio was calculated as the proportion of tdT+ cells out of total DAPI-positive epidermal cells per image. Proliferation and death rates were expressed as the proportion of EdU-positive or CC3-positive cells out of total DAPI-positive cells, either across the whole epidermis or within each tdT-positive and tdT-negative population separately. Cell-intrinsic rates were calculated by normalising EdU-positive or CC3-positive counts within each population to the total number of cells of the same genotype (e.g. EdU+,tdT+/total tdT+ cells).

### Flow cytometry

#### Mouse epidermal keratinocyte isolation and staining

Single-cell suspensions were prepared from mouse epidermis following an established protocol^23^. Briefly, skin was dissected and placed dermis-side down onto Whatman paper in a 6-well plate containing dispase solution (1:10 in 1XPBS). Plates were incubated overnight at 4°C on a shaker to allow for epidermal-dermal separation. The following day, the epidermis was peeled away from the dermis, cut into pieces, and incubated in 0.25% trypsin at 37°C for up to 15 minutes to dissociate cells. Trypsin activity was quenched by addition of ice-cold 1XPBS supplemented with 2% FBS, and the suspension was pipetted vigorously to release single cells. The cell suspension was filtered through a 70 µm cell strainer into a 50 ml Falcon tube, washed twice with 2% FBS in PBS, and centrifuged at 300 g for 5 minutes at 4°C. The resulting pellet was resuspended in FACS buffer for downstream staining.

For flow cytometric analysis of X-skewing ratios, single-cell suspensions were stained with a lineage exclusion panel comprising biotinylated antibodies against CD31, CD45, CD117, and CD140a to exclude non-keratinocyte populations, alongside PE-Cy7-conjugated CD49f to identify epidermal keratinocytes. tdTomato fluorescence was used to distinguish wild-type (tdT-positive) from mutant (tdT-negative) cells within the keratinocyte gate. Flow cytometry was performed using the ZE5 Cell Analyzer (Bio-Rad) and data were analysed using FlowJo™ v10.8 Software (BD Life Sciences).

### Statistical analyses

Statistical analyses were performed using GraphPad Prism (version 11.0.0 for macOS, GraphPad Software, Boston, Massachusetts USA, www.graphpad.com). The choice of statistical test was determined by data distribution, assessed using the Shapiro-Wilk normality test for smaller datasets and the D’Agostino-Pearson for larger ones.

For comparisons between two independent groups with continuous normally distributed data, unpaired t-tests with Welch’s correction were used to account for unequal variances between these groups. For data that were not distributed normally or data with many zero values, non-parametric alternatives were used: Mann-Whitney test for unpaired comparisons and Wilcoxon matched-pairs signed rank test for paired comparisons within the same tissue section or animal. For comparisons where one group had zero variance (e.g., wild-type cell death rates), one-sample t-tests and Wilcoxon tests against a theoretical mean of zero were used.

For analyses across multiple timepoints shown in Figure 1F (left panel), two-way ANOVA was used, with genotype and timepoint as the two factors and their interaction term included in the model. Post-hoc pairwise comparisons were performed using Tukey’s multiple comparisons test. For within-genotype analyses (centre and right panels), one-way ANOVA was used, followed by Tukey’s multiple comparisons test.

For the two-way ANOVAs in Fig.2, pre-specified comparisons between genotypes at each timepoint were performed using Fisher’s least significant difference (LSD) test without correction for multiple comparisons. At P3, the relationship between local tissue composition and proliferation was assessed by fitting a binomial generalised linear mixed model (using the lme4 package (v1.1-35.1) in R (v4.3.3)) to field-level counts of EdU⁺ cells among tdT⁺ basal cells in *Nemo +/-* animals (71 fields from 4 animals), with the proportion of tdT⁺ cells per field as a fixed effect, a random intercept for animal to account for the nesting of fields within animals, and a field-level random intercept to account for overdispersion. Significance of the fixed effect was assessed by a Wald test. An equivalent model was fitted independently to the tdT⁻ population.

Graphs were plotted using GraphPad Prism (version 11.0.0). Data are presented as individual data points with mean or median indicated where appropriate.

### scRNA sequencing library preparation

Single-cell RNA sequencing was performed using the Chromium Single Cell Gene Expression Flex technology from 10x Genomics. In this experiment, 16 epidermal samples from mice at different timepoints ( E17.5, P0, P3, P7) and conditions (*Nemo +/-*,WT) were collected. During library preparation, three custom probe pairs targeting different regions of the tdTomato gene were incorporated (see table). These probes, barcoded as BC001-BC016, enabled the pooling and sequencing of all samples in a single run.

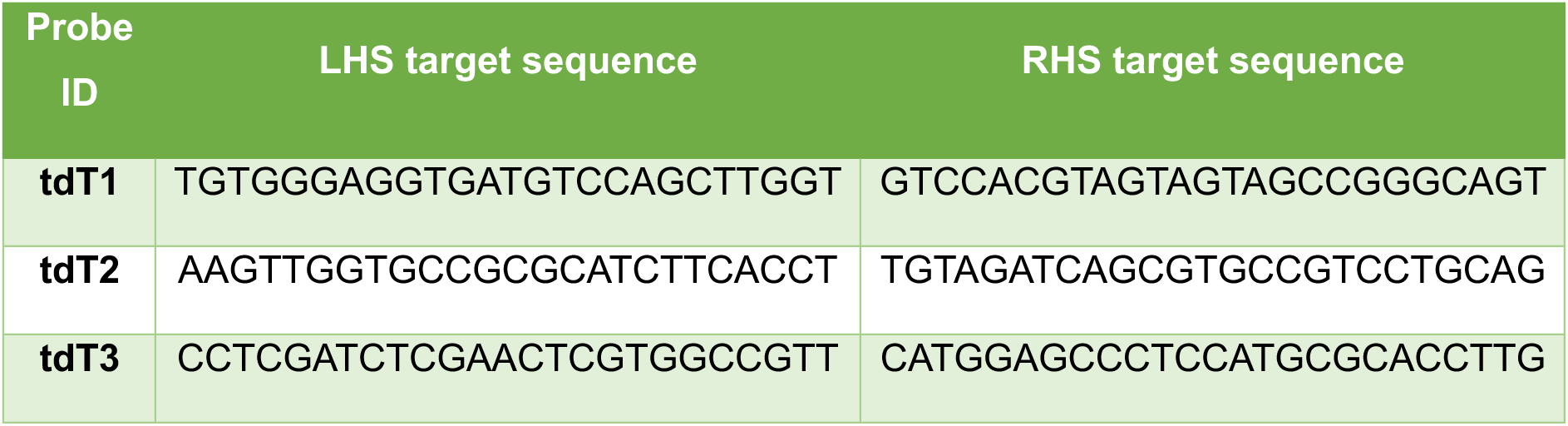

Raw data were pre-processed using Cell Ranger version 9.0.0 using the command “multi” to simultaneously process the transcriptome and demultiplex the data. During this step, the Probe Set Reference CSV file “Chromium_Mouse_Transcriptome_Probe_Set_v1.1.1_GRCm39-2024-A.csv” was modified to include the custom probe sequences, and alignment was performed using the mouse reference genome assembly “refdata-gex-GRCm39-2024-A”.

Samples were processed using R (version 4.1.0) and Seurat (version 5.0.1)^24^. Ambient RNA was removed using DecontX^25^ and doublets were identified and eliminated using scDblFinder^26^. Quality assessment criteria retained only cells expressing a minimum of 500 genes, with at least 500 reads, and limited mitochondrial gene expression to a maximum of 7% for subsequent analyses.

Afterwards, samples were normalised and scaled regressing against mitochondrial gene percentages variables to mitigate their influence as confounding sources of variation. For dimensionality reduction, 30 dimensions were utilised in invoking Seurat functions including RunPCA, FindNeighbors, and RunUMAP. Subsequently, samples were integrated using the FastMNN method^27^. Clustering was performed utilizing the Leiden algorithm on the shared nearest neighbour graph prior to cell annotation, as suggested by Heumos, et al.^28^

Cell type identity was assigned based on canonical marker gene expression, featuring 12 major populations, namely basal, suprabasal, wound, stress keratinocytes, hair follicle progenitors, hair shaft cells, immune niche cells, melanocytes, and stromal populations. Wound and stress keratinocytes were defined by expression of stress keratins (Krt6a, Krt16, Krt17), AP-1 transcription factors, and matrix remodelling genes, and were identified exclusively in the Nemo^+/−^ epidermis (see **Supplemental Fig. 3**).

### scRNA sequencing data analysis

#### Compositional analysis

Cell type proportions were calculated per sample at each timepoint. Statistically significant differences in cell type composition between Nemo^+/-^ and WT animals were identified using Fisher’s exact test, with Benjamini-Hochberg correction for multiple comparisons. X-skewing ratios were quantified as the proportion of tdT-positive cells over total cells within the basal keratinocyte population per animal per timepoint.

#### Differential expression analysis

Differential expression analysis comparing tdT-positive versus tdT-negative cells within the wound cluster at P3. Analysis was performed using Seurat FindMarkers function^29^ with the Wilcoxon rank-sum test and Bonferroni correction for multiple testing (FDR<0.05). A minimum log2 fold change threshold of 0.10 and minimum cell fraction threshold of 0.1 were applied. Reporter transgene sequences (Hprt, TDT1–3) were excluded from all comparisons prior to ranking to avoid technical artefacts. Results were visualised as volcano plots using ggplot2^30^.

#### Gene set enrichment analysis

Differentially expressed genes were converted to Entrez IDs using the org.Mm.eg.db^31^ annotation database. Over-representation analysis (ORA) was performed separately for genes upregulated in tdT-positve and tdT-negative cells using the clusterProfiler enrichKEGG function for KEGG pathway enrichment and enrichGO for Gene Ontology Biological Process terms, with organism code “mmu”, p-value cut-off of 0.05, and Benjamini-Hochberg correction for multiple testing. Results were visualised as dotplots showing gene ratio and adjusted p-value.

#### Pseudotime trajectory analysis

Differentiation trajectories were reconstructed using Monocle3^32^. The MNN-corrected UMAP embedding from Seurat was imported directly into Monocle3 as a CellDataSet object containing approximately 49,000 keratinocytes across all postnatal timepoints, plus the E17.5 timepoint used for root placement. The principal graph was learned on the UMAP embedding using the learn_graph function, and pseudotime values were computed using order_cells with the root node placed in the E17.5 basal keratinocyte cluster (6,188 root cells).

#### Cell-cell communication analysis

Cell-to-cell signalling was inferred using CellChat (version 1)^33^. To examine TNF signalling from immune to epithelial populations, CellChat objects were reconstructed for WT and Nemo+/- epidermis at P3, using all annotated cell types as input groups. Ligand-receptor interaction probabilities were computed using the CellChat mouse signalling database with a minimum cell threshold 10 cells and triMean aggregation of gene expression values. Communication networks were computed using computeCommunProb and aggregated at the pathway level using computeCommunProbPathway. TNF pathway signalling was visualised using circular network plots and bubble plots showing communication probability per receiver cell type. Incoming signalling strength per cell type was defined as the column sum of the communication probability weight matrix.

To examine signalling between tdT-positive and tdT-negative epidermal populations, a second CellChat analysis was performed on the Nemo +/- epidermis pooled across timepoints and in parallel on the WT epidermis. Cell identities were defined as the combination of cell type annotation and TDT status (e.g. “Wound keratinocytes (wt)”, “Wound keratinocytes (mut)”), yielded 23 distinct groups in Nemo +/- and 20 in WT, each with a minimum of 30 cells. Disease-specific signalling pathways were defined as those detected in the Nemo +/- network but absent from the WT baseline.

#### Pseudobulk DESeq2

To test for differences in EGF ligand and receptor expression (*Epgn, Hbegf, Tgfa, Areg, Egfr, Erbb2, Erbb3*) between tdT-positive and tdT-negative cells, raw counts were aggregated per animal per TDT group within wound, stress, and basal keratinocyte clusters across timepoints, yielding 6 pseudobulk samples per group (n=2 animals per timepoint). Differential expression was performed using DESeq2 with TDT status as the design factor, retaining genes with counts ≥5 in at least 2 samples. Adjusted p-values were calculated using Benjamini-Hochberg correction across the 7 genes tested.

#### EGF expression analysis

Normalised expression values for EGF ligands (*Epgn, Hbegf, Tgfa, Areg)* and receptors (*Egfr, Erbb2, Erbb3)* were extracted from the integrated Seurat object and stratified by TDT status and timepoint within wound, stress, and basal keratinocyte clusters in Nemo +/- animals. Percentage of expressing cells and mean expression per group were visualised as violin plots. Per-timepoint log2 ratios of mean expression (tdT-/tdT+) were calculated descriptively and displayed as heatmap.

#### Cluster marker dotplot

Cluster transcriptional identity was summarised using Seurat’s DotPlot function across all cells, genotypes, and timepoints, displaying five to six canonical marker genes per cluster.

## Data and code availability

The single-cell RNA-sequencing data generated in this study will be deposited in the Gene Expression Omnibus (GEO) at journal submission and made publicly available upon journal publication. Accession numbers will be provided at that time. All other data supporting the findings of this study are available from the corresponding author upon reasonable request. Analysis code is available from the corresponding author upon request and will be deposited upon publication.

## Contributions

S.J.E. conceived and supervised the project, and wrote the manuscript. E.A. performed most experiments, analysis and wrote the manuscript. K.G. assisted with flow cytometry in Fig. 1 and performed qPCR experiments. M.v.W. and V.S. helped with image analysis. L.S. performed analysis of immune depletion. I.O. and J.G. performed scRNA-seq raw data processing, quality control and sample integration. I.F. assisted with histology and cryosectioning. S.N. provided technical assistance throughout the project.

## Supplemental Figure Legends

**Supplemental Figure S1.**
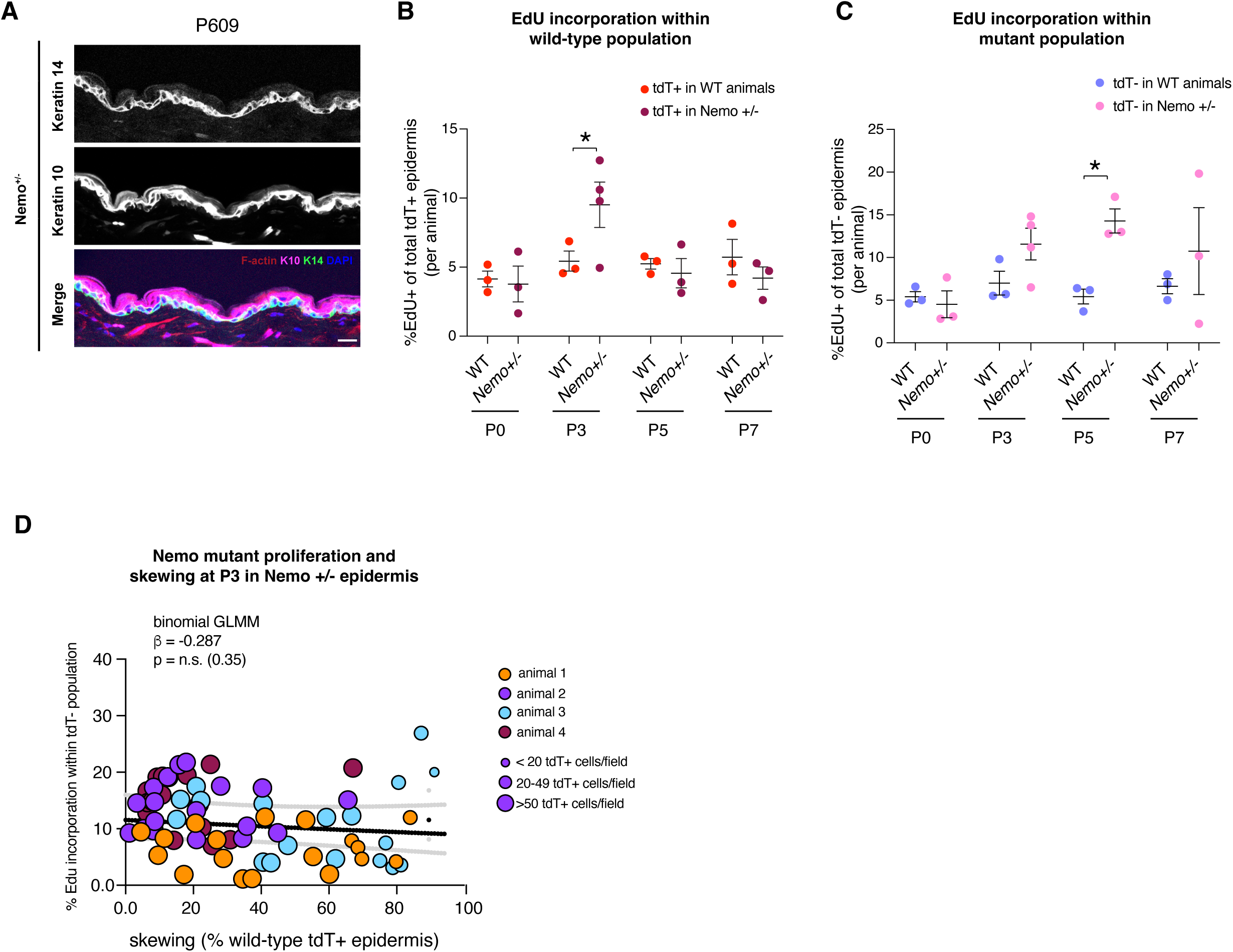
Long-term recovery of Nemo +/- epidermis and proliferation in the wild-type and mutant populations. **(A)** Representative immunofluorescence of *Nemo +/-* back skin at P609 stained for Keratin 14 (K14, basal), Keratin 10 (K10, suprabasal), F-actin and DAPI, showing restored epidermal architecture. Scale bar, 20 µm. (**B**, **C**) Percentage of EdU+ cells within the tdT+ (B) and tdT– (C) epidermal populations of wild-type and *Nemo +/-* animals at P0, P3, P5 and P7 (1 h EdU pulse). Each dot represents one animal (n = 3–4 per genotype and timepoint); bars show mean ± SEM; two-way ANOVA (genotype × timepoint) on animal means followed by uncorrected Fisher’s LSD tests comparing genotypes at each timepoint. (D) EdU incorporation in the tdT– (mutant) population as a function of local skewing (fraction of tdT+ basal cells) per field of view in P3 Nemo +/- epidermis, analysed as in Fig. 2H. Each point is one field; colour indicates animal (n = 4 animals) and point size the number of tdT– cells per field. Line shows a binomial generalised linear mixed model fit (random intercepts for animal and field; n = 71 fields from 4 animals), with 95% confidence band. No significant relationship was detected (β = −0.29 log-odds per unit fraction, P = 0.35). *P < 0.05.

**Supplemental Figure S2.**
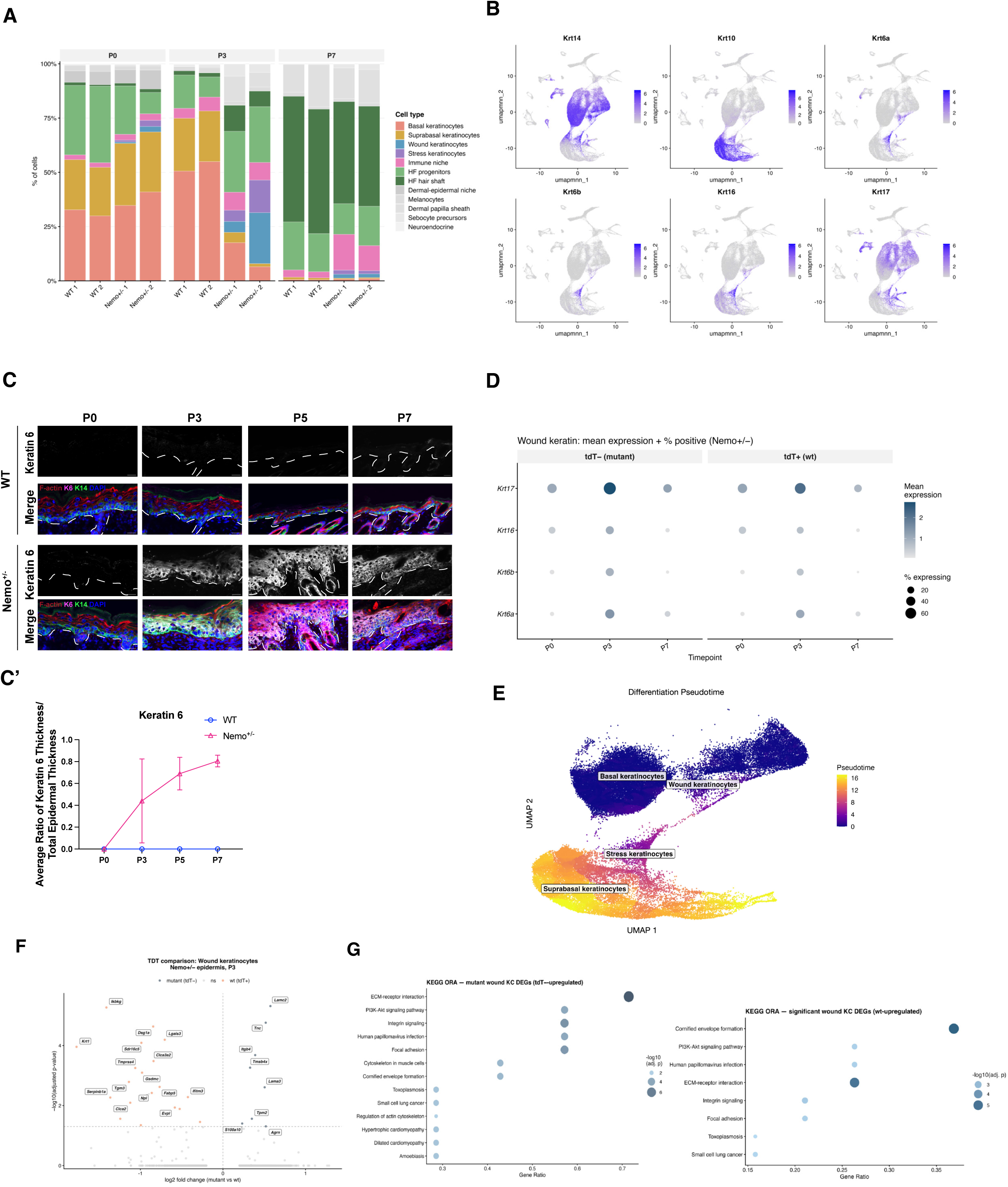
Wound-like keratinocytes expand in *Nemo +/-* epidermis and are enriched for mutant cells. **(A)** Cell-type composition of each scRNA-seq sample (WT 1–2, *Nemo +/-* 1–2) at P0, P3 and P7. **(B)** Feature plots of *Krt14*, *Krt10* and the wound-associated keratins *Krt6a*, *Krt6b*, *Krt16* and *Krt17* on the integrated UMAP. **(C)** Immunofluorescence for Keratin 6 (K6) with K14, F-actin and DAPI in wild-type and *Nemo +/-* back skin at P0, P3, P5 and P7. Dashed line marks the basement membrane. Scale bars, 20 µm. **(C′)** Ratio of K6+ thickness to total epidermal thickness over time (n = 1 animal per genotype and timepoint; mean ± [SD]). **(D)** Mean expression (colour) and percentage of expressing cells (dot size) of wound keratins in tdT– (mutant) and tdT+ (wild-type) cells of *Nemo +/-* epidermis at P0, P3 and P7. **(E)** Differentiation pseudotime (Monacle) across basal, wound, stress and suprabasal keratinocytes. **(F)** Volcano plot of differentially expressed genes between tdT– (mutant) and tdT+ (wild-type) wound keratinocytes in P3 *Nemo +/-* epidermis (n = 2 animals pooled). Cell-level Wilcoxon rank-sum test (Seurat FindMarkers; minimum log2 fold change 0.1, minimum detection fraction 0.1); dashed line indicates Bonferroni-adjusted *P* = 0.05. Reporter transgene sequences were excluded before testing. **(G)** KEGG over-representation analysis of genes upregulated in mutant (tdT–, left) and wild-type (tdT+, right) wound keratinocytes. Dot size, –log10 adjusted *P*.

**Supplemental Figure S3.**
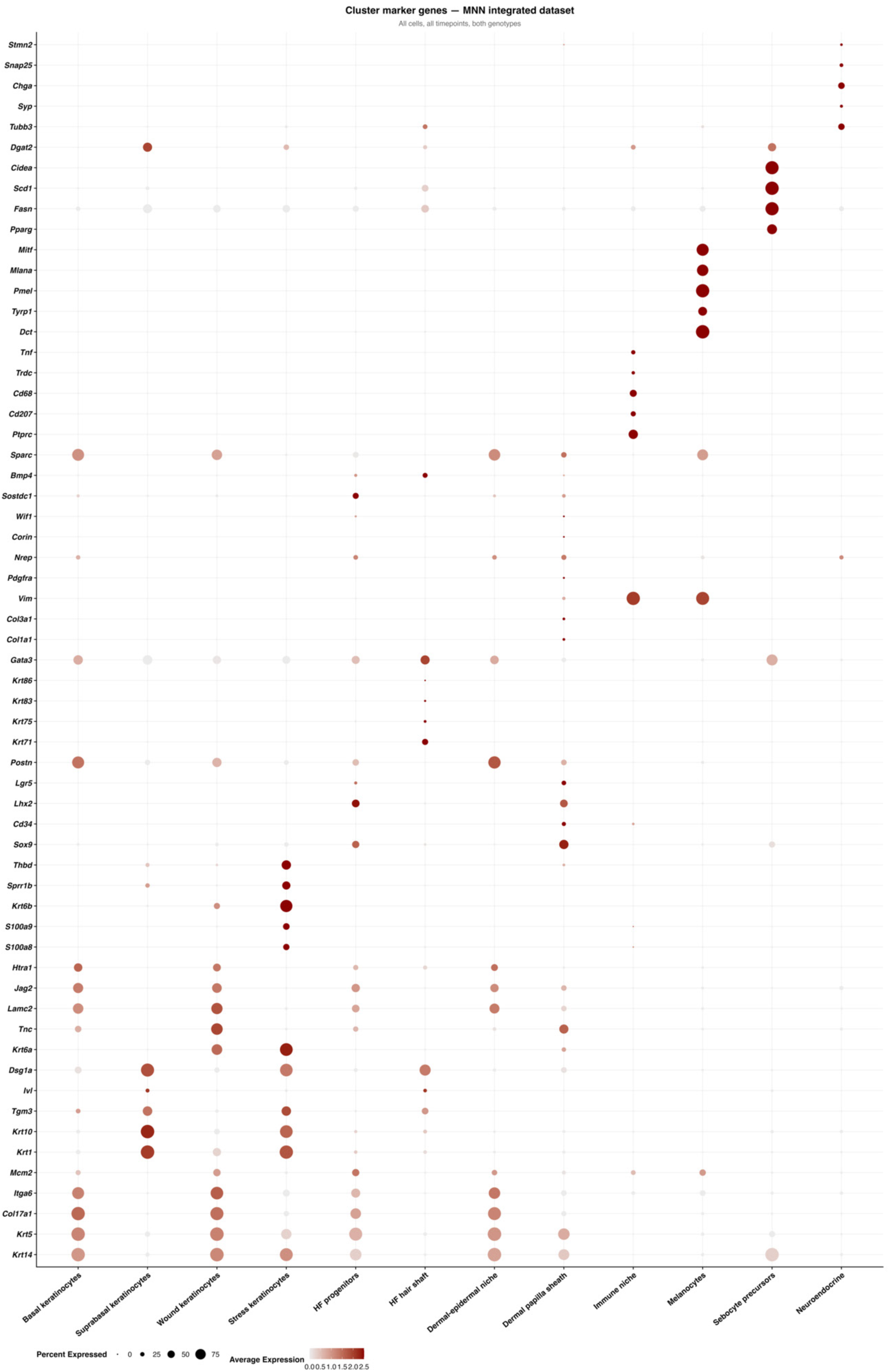
Cluster marker genes. Dot plot of marker gene expression across annotated cell types in the MNN-integrated scRNA-seq dataset (all cells, all timepoints, both genotypes). Colour indicates average scaled expression and dot size the percentage of cells expressing each gene.

**Supplemental Figure S4.**
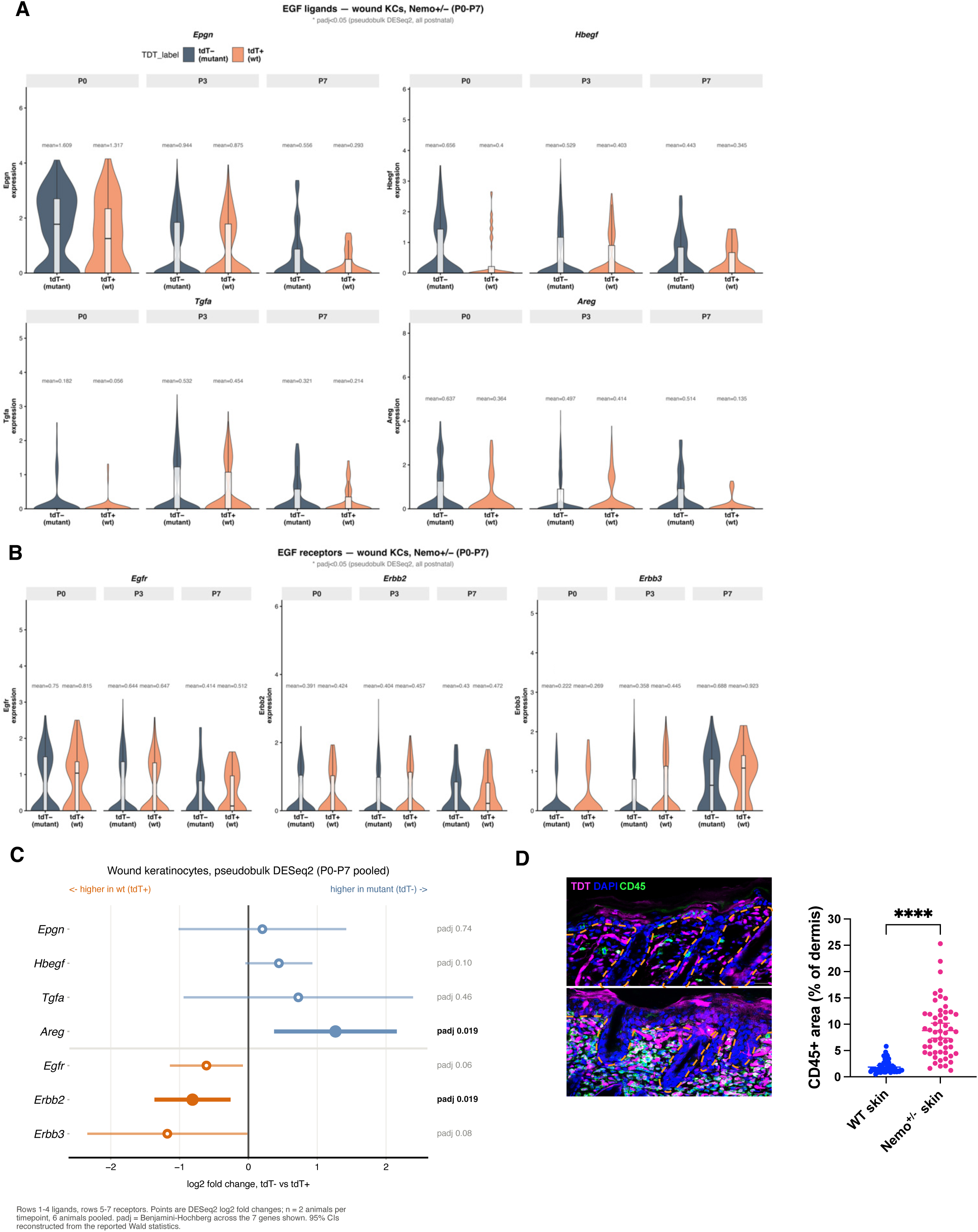
EGF-family ligand–receptor asymmetry between mutant and wild-type wound keratinocytes, and immune infiltration in *Nemo +/-* skin. **(A, B)** Violin plots of EGF-family ligand (A; *Epgn*, *Hbegf*, *Tgfa*, *Areg*) and receptor (B; *Egfr*, *Erbb2*, *Erbb3*) expression in tdT– (mutant) and tdT+ (wild-type) wound keratinocytes of *Nemo +/-* epidermis at P0, P3 and P7; mean expression is indicated above each violin. **(C)** Pseudobulk DESeq2 log2 fold changes (tdT– vs tdT+) for the genes in (A, B), with P0–P7 pooled (n = 2 animals per timepoint, 6 animals in total). Error bars show 95% confidence intervals reconstructed from the Wald statistics; adjusted *P* values (Benjamini–Hochberg across the seven genes) are shown on the right. *Areg* is significantly higher in mutant cells and *Erbb2* in wild-type cells (adjusted *P* = 0.019 each). **(D)** Representative images and quantification of CD45+ area (% of dermis) in wild-type and *Nemo +/-* skin at P5, stained for CD45, tdTomato and DAPI. Each dot represents one field of view from n = 3 animals per genotype; mean ± SEM; t-test. \*\*\*\**P* < 0.0001.

